# Identification of antiviral drug candidates against SARS-CoV-2 from FDA-approved drugs

**DOI:** 10.1101/2020.03.20.999730

**Authors:** Sangeun Jeon, Meehyun Ko, Jihye Lee, Inhee Choi, Soo Young Byun, Soonju Park, David Shum, Seungtaek Kim

## Abstract

COVID-19 is an emerging infectious disease and was recently declared as a pandemic by WHO. Currently, there is no vaccine or therapeutic available for this disease. Drug repositioning represents the only feasible option to address this global challenge and a panel of 48 FDA-approved drugs that have been pre-selected by an assay of SARS-CoV was screened to identify potential antiviral drug candidates against SARS-CoV-2 infection. We found a total of 24 drugs which exhibited antiviral efficacy (0.1 μM < IC_50_ < 10 μM) against SARS-CoV-2. In particular, two FDA-approved drugs - niclosamide and ciclesonide – were notable in some respects. These drugs will be tested in an appropriate animal model for their antiviral activities. In near future, these already FDA-approved drugs could be further developed following clinical trials in order to provide additional therapeutic options for patients with COVID-19.

## Introduction

COVID-19 is an emerging infectious disease caused by a novel coronavirus, SARS-CoV-2 ^1^. Although the case fatality rate due to this viral infection varies from 1 to 12% ^2^, the transmission rate is relatively high ^3^ and recently, the WHO declared COVID-19 outbreak a pandemic. Currently, there is no vaccines or therapeutics available and the patients with COVID-19 are being treated with supportive care.

Drug repositioning could be an effective strategy to respond immediately to emerging infectious diseases since the new drug development usually takes more than 10 years ^4^. FDA-approved drugs provide safe alternatives only in the case where at least modest antiviral activity can be achieved. Accordingly, several drugs are being tested in numerous clinical trials ^5^ including remdesivir, lopinavir, and chloroquine ^6^.

In this study, we screened a panel of FDA-approved drugs to identify antiviral drug candidates for the treatment of COVID-19 and suggest the identified drug candidates may be considered for therapeutic development.

## Results and Discussion

We screened approximately 3,000 FDA- and IND-approved drug library against SARS-CoV to identify antiviral drug candidates (manuscript in preparation). Since the SARS-CoV and SARS-CoV-2 are very similar (79.5% sequence identity) ^1^, the drugs which show antiviral activity against SARS-CoV are expected to show similar extent of antiviral activity against SARS-CoV-2.

A total of 35 drugs were selected from the earlier SARS-CoV screening results. In addition, 13 drugs were included based on recommendations from infectious diseases specialists (Table 1). For screening experiments, Vero cells were used and each drug was added to the cells prior to the virus infection. At 24 h after the infection, the infected cells were scored by immunofluorescence analysis with an antibody specific for the viral N protein of SARS-CoV-2. The confocal microscope images of both viral N protein and cell nuclei were analyzed using our in-house Image Mining (IM) software and the dose-response curve (DRC) for each drug was generated (Figure 1).

**Table 1.**
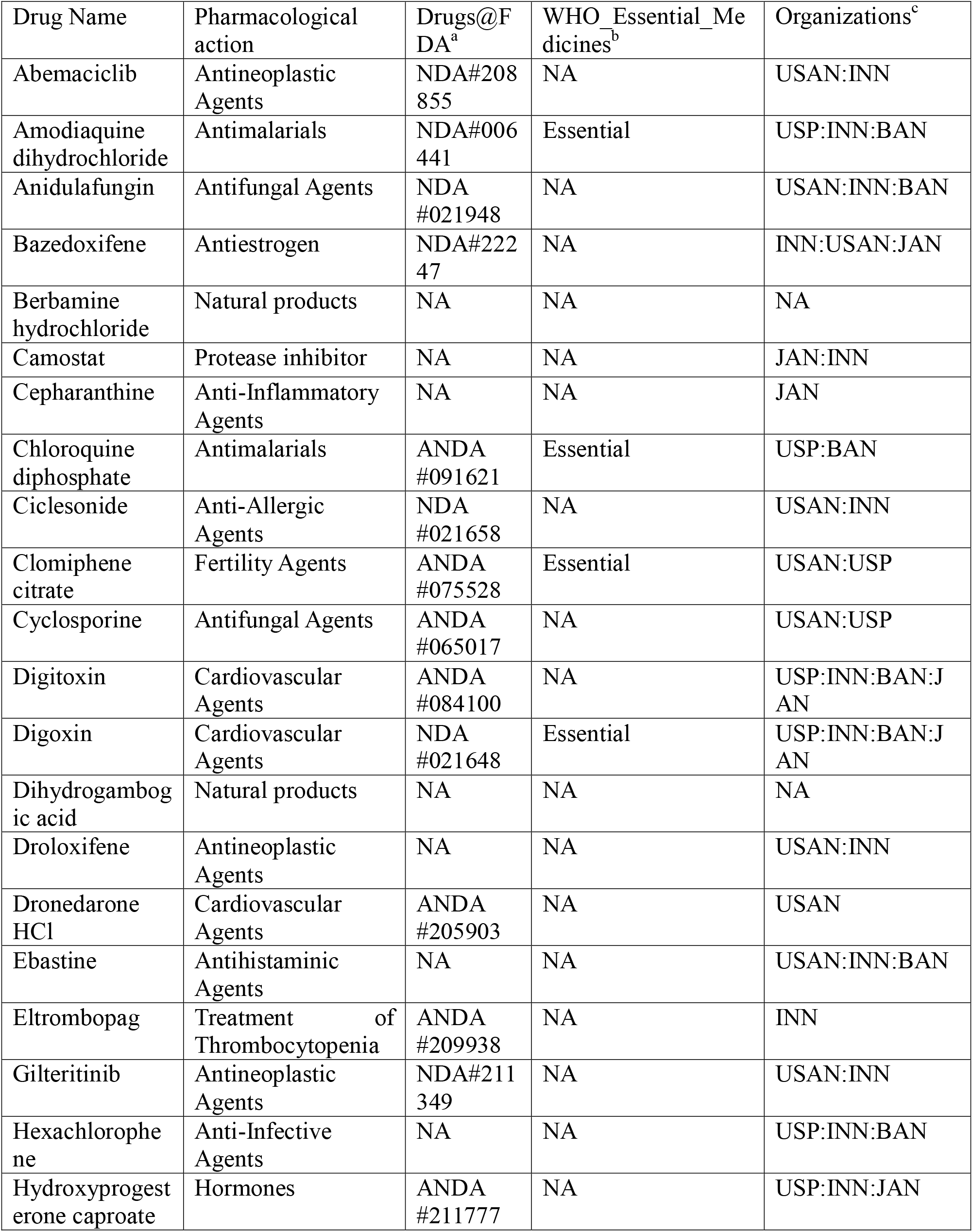
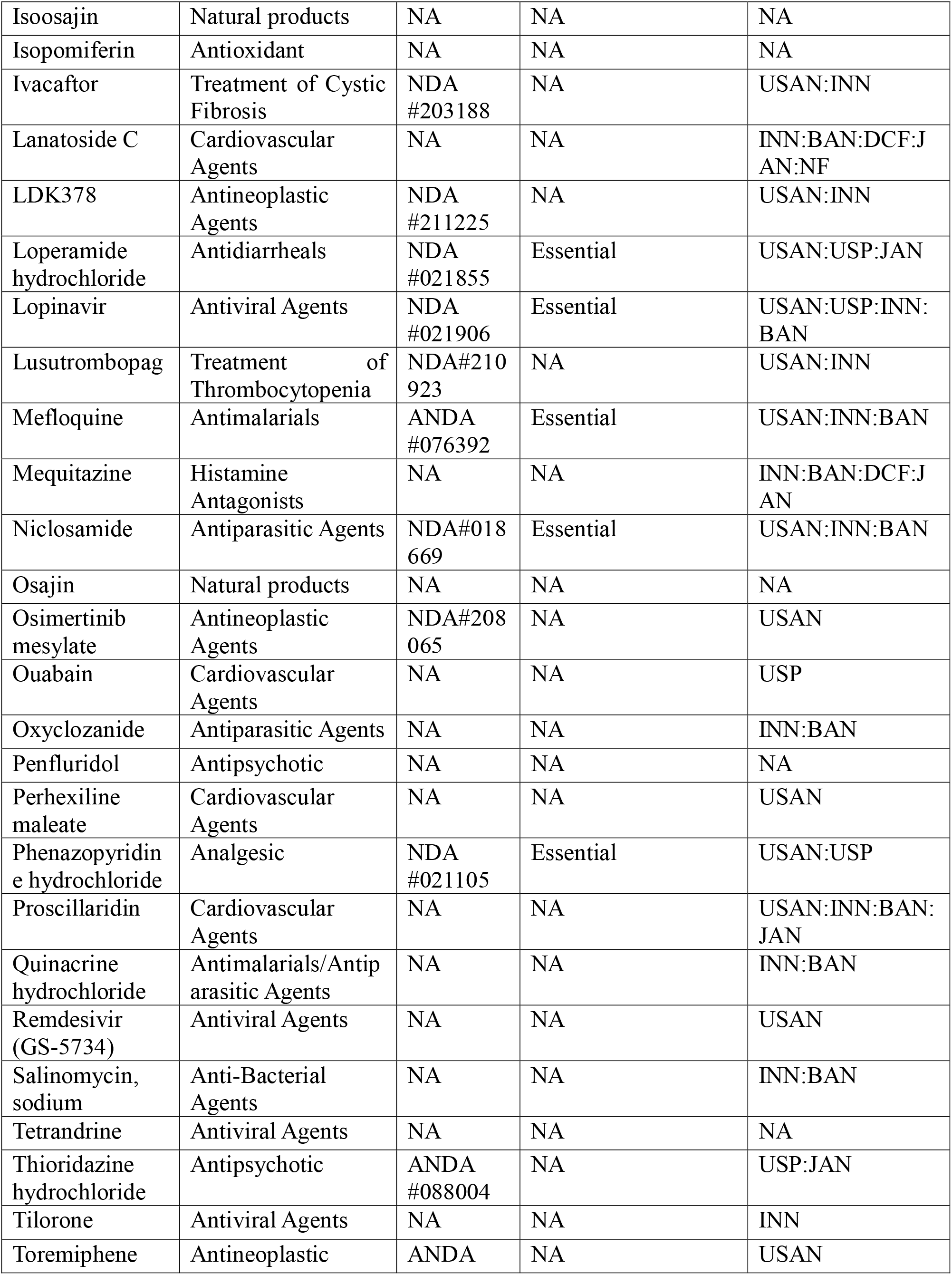
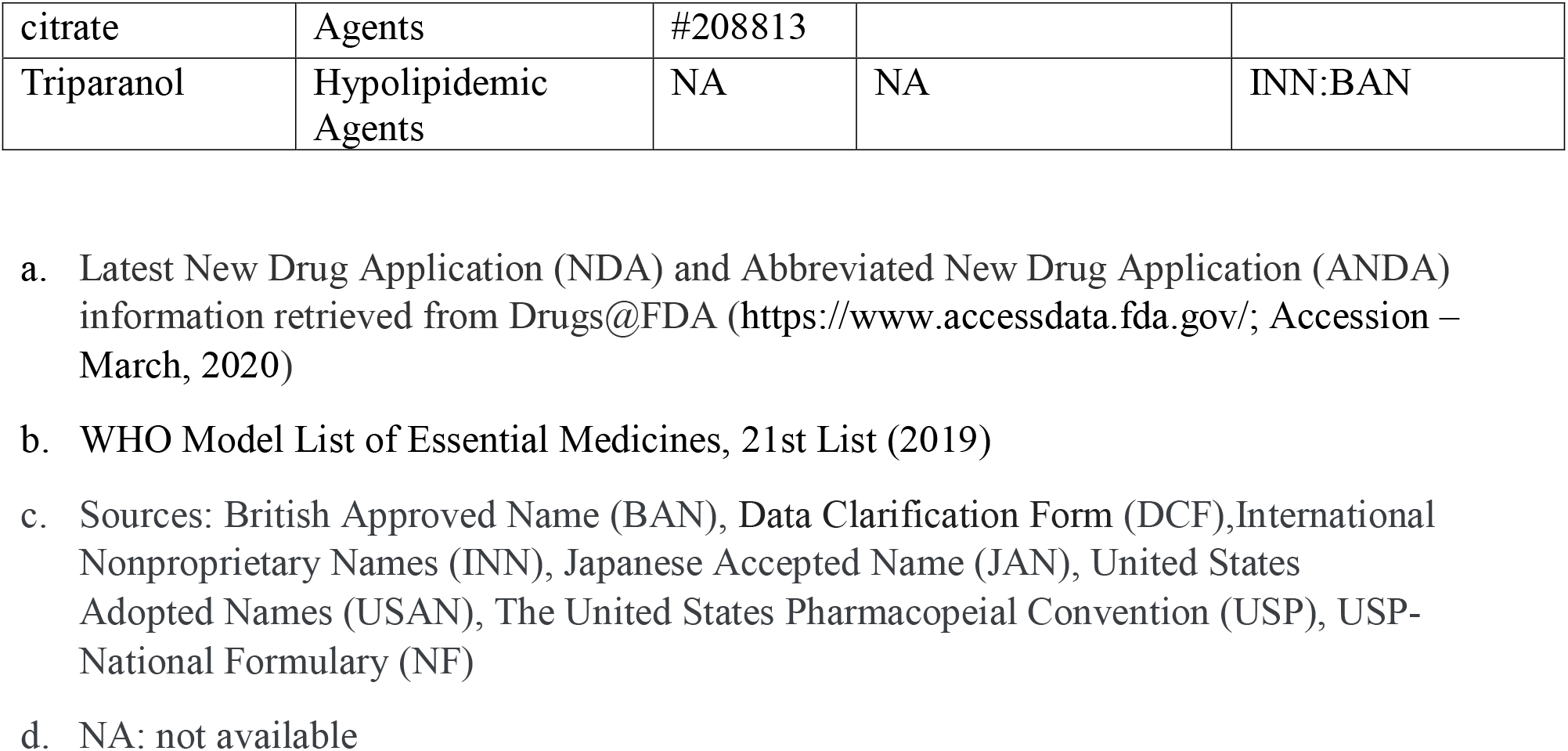
Pharmacological actions and registration status of drugs

**Figure 1.**
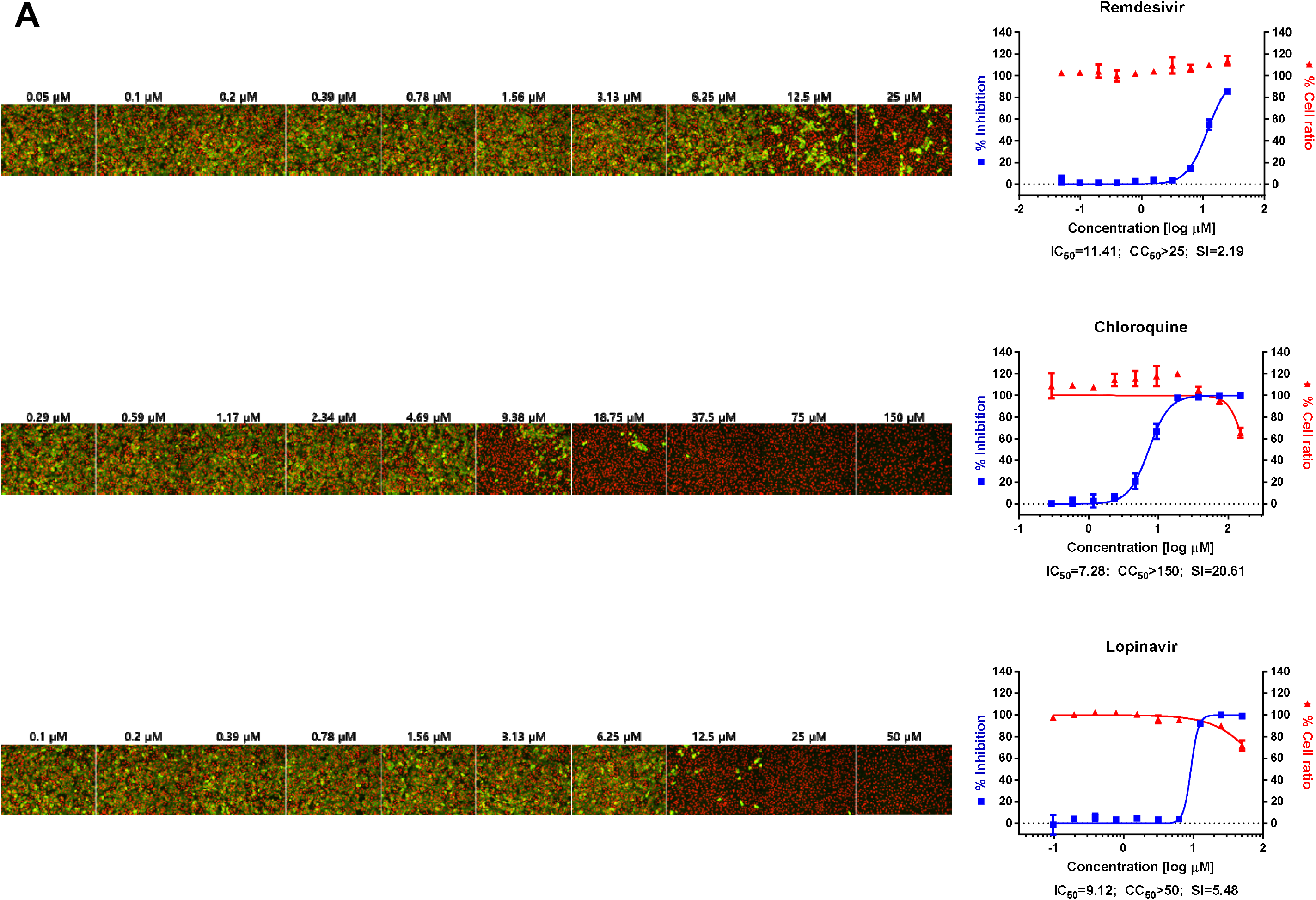
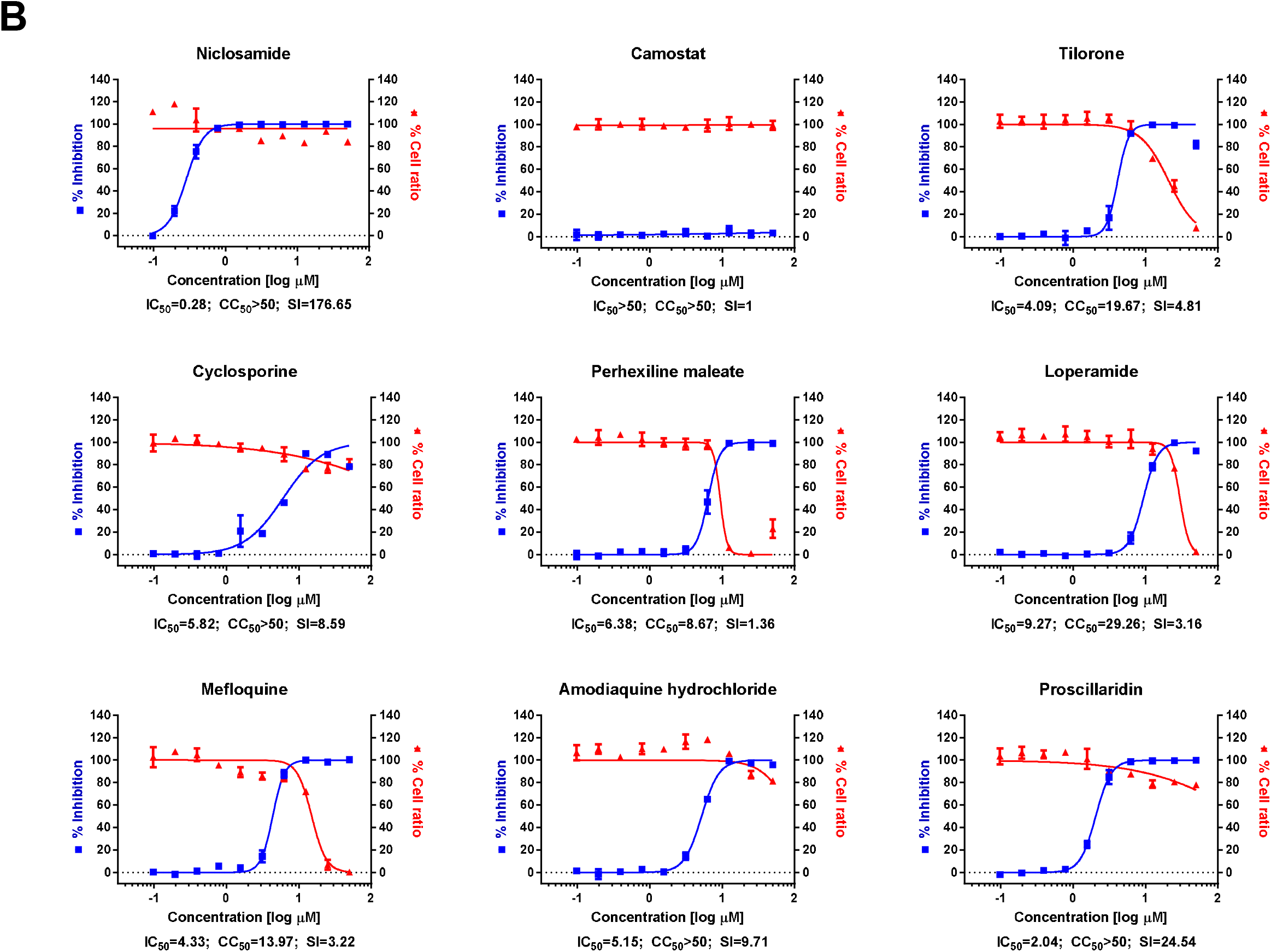
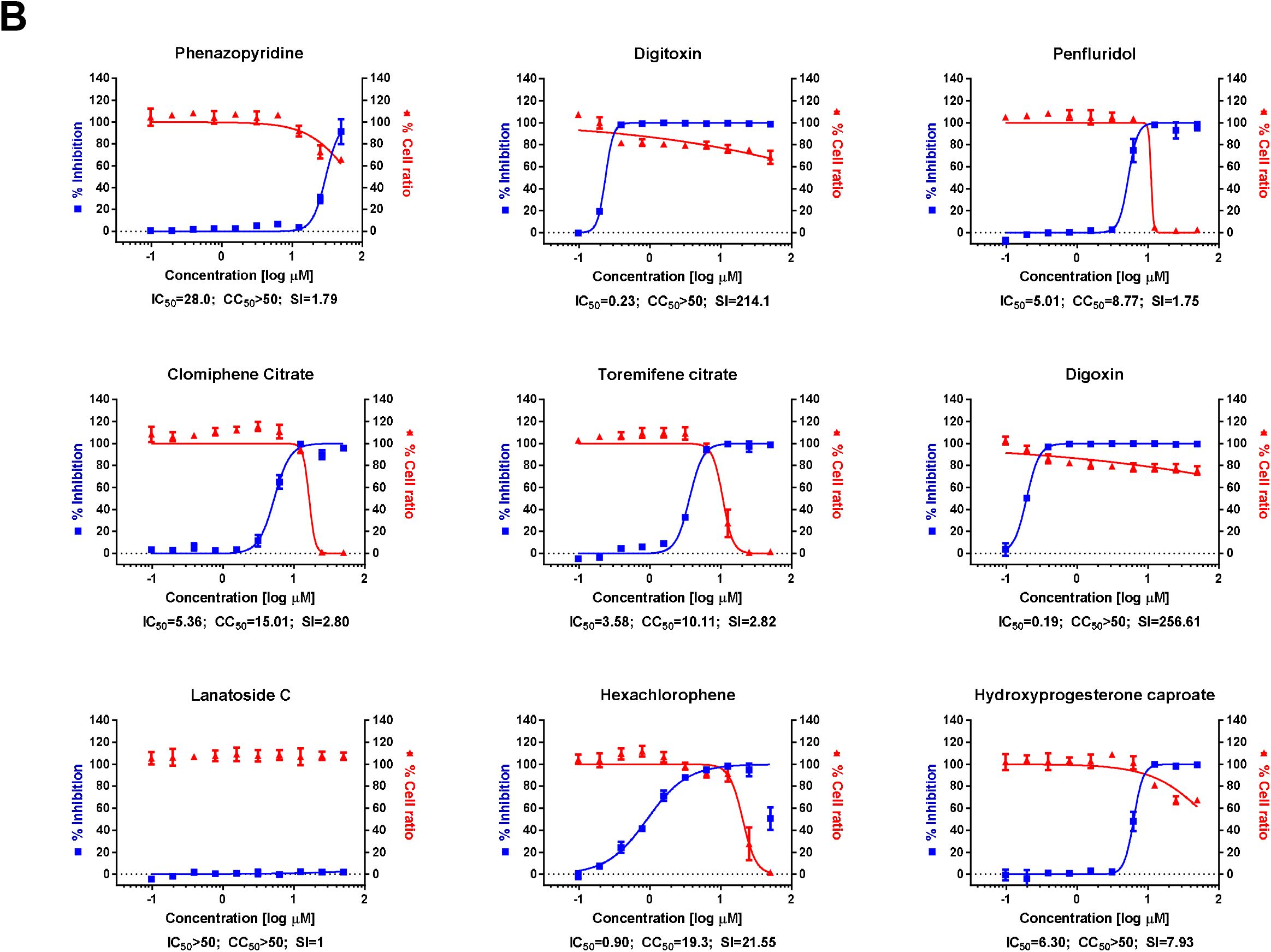
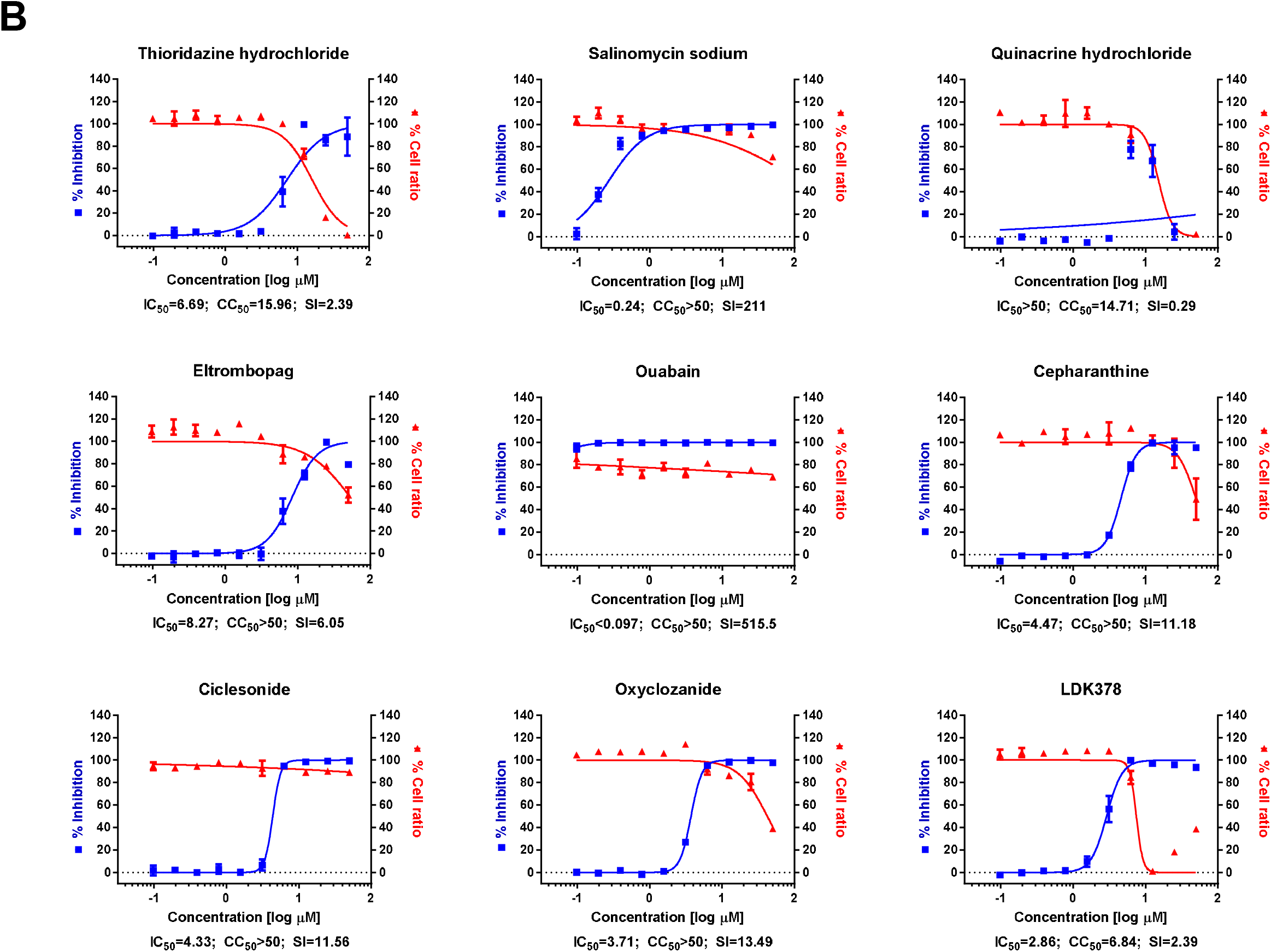
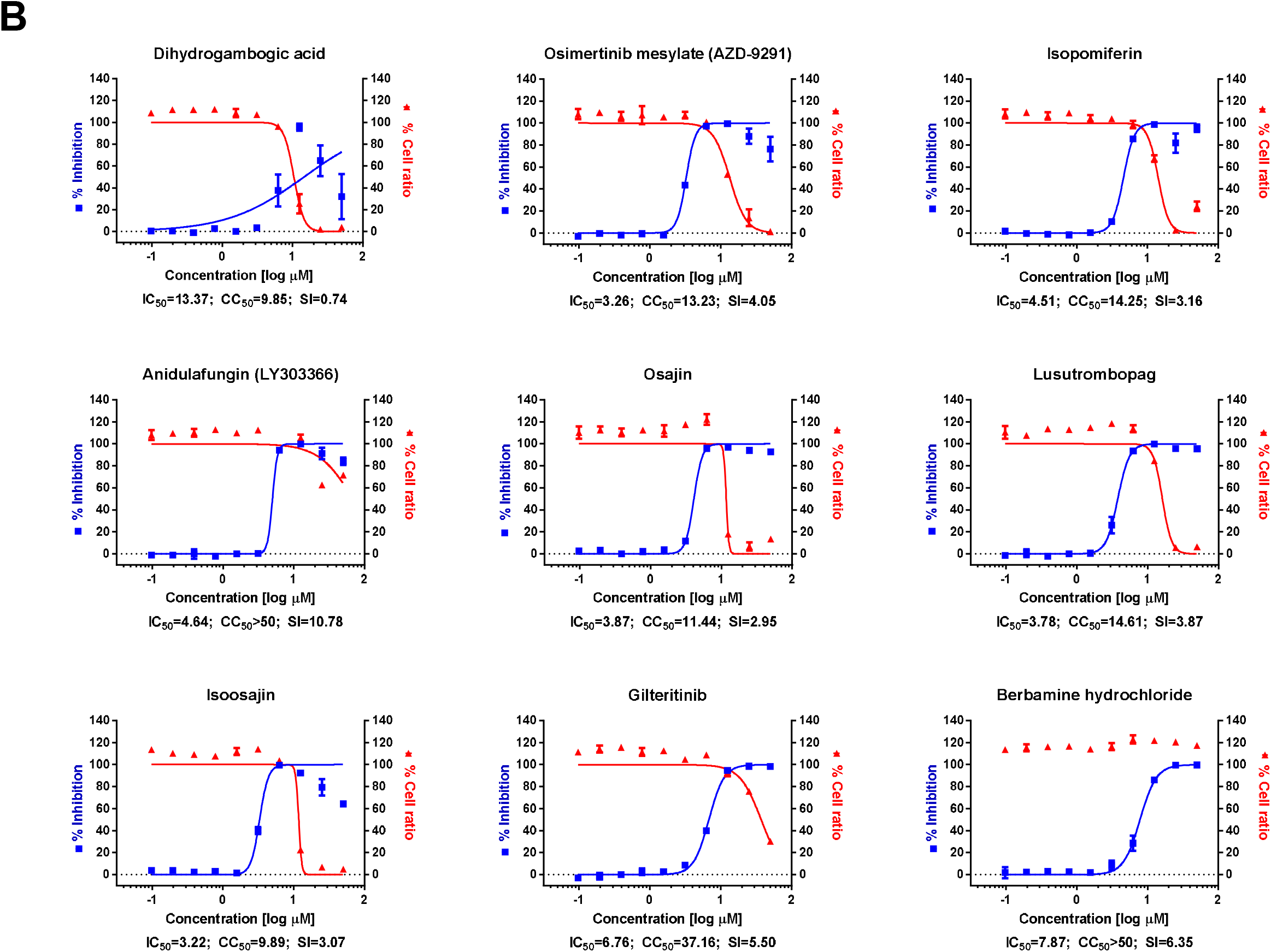
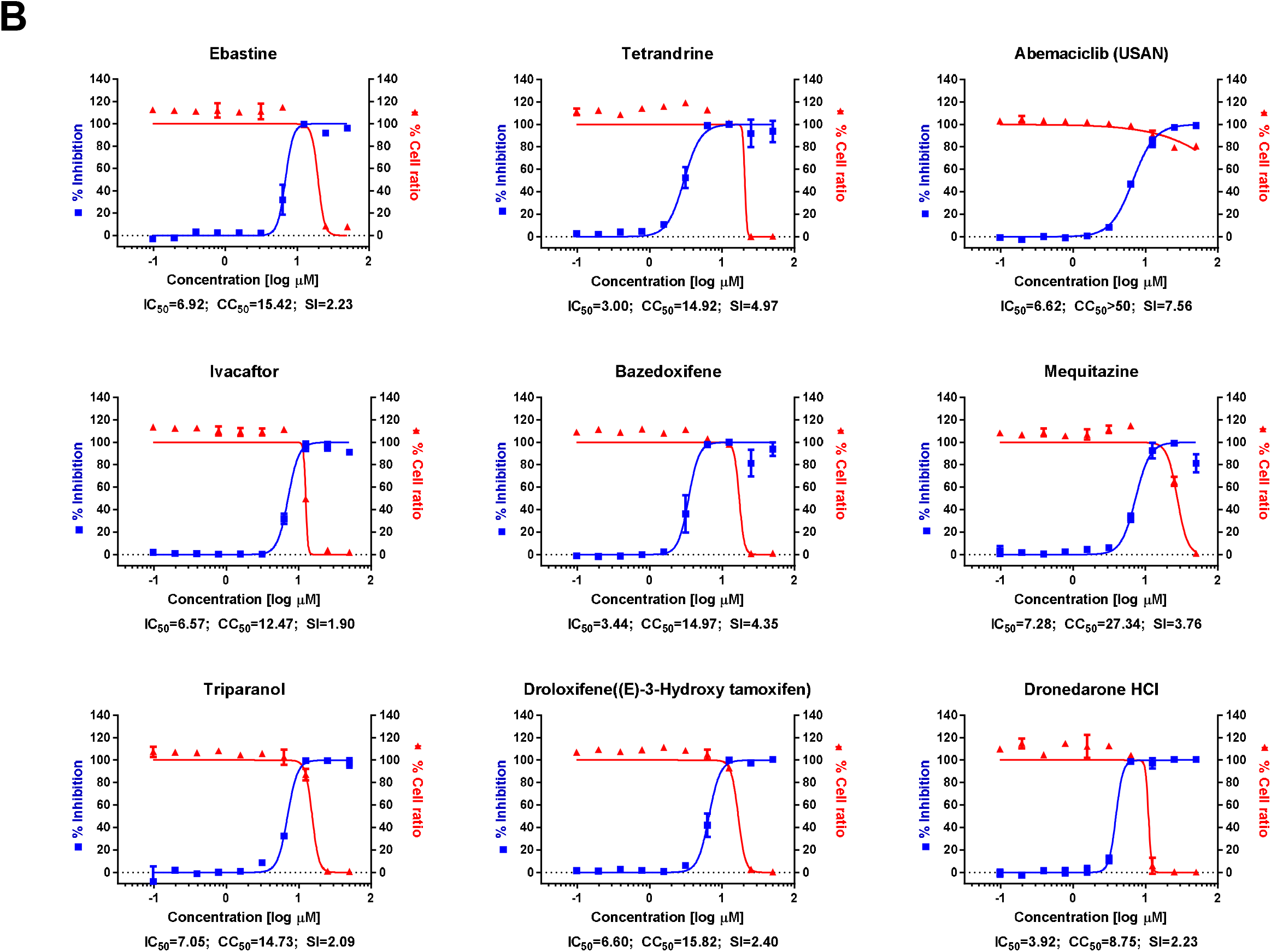
(A) Dose-response curve analysis by immunofluorescence for reference drugs. The blue squares represent inhibition of virus infection (%) and the red triangles represent cell viability (%). The confocal microscope images show cell nuclei (red) and viral N protein (green) at each drug concentration. Means ± SD were calculated from duplicate experiments. (B) Dose-response curve analysis by immunofluorescence for 45 drugs that were tested in this study. The blue squares represent inhibition of virus infection (%) and the red triangles represent cell viability (%). Means ± SD were calculated from duplicate experiments.

Chloroquine, lopinavir, and remdesivir were used as reference drugs with IC_50_ values of 9.12, 7.28, and 11.41 μM, respectively (Figure 1A). Among the 48 drugs that were evaluated in our study, 24 drugs showed potential antiviral activities against SARS-CoV-2 with IC_50_ values in between 0.1 and 10 μM; Tilorone, Cyclosporine, Loperamide, Mefloquine, Amodiaquine, Proscillaridin, Digitoxin, Digoxin, Hexachlorophene, Hydroxyprogesterone caproate, Salinomycin, Ouabain, Cepharanthine, Ciclesonide, Oxyclozanide, Anidulafungin, Gilteritinib, Berbamine, Tetrandrine, Abemaciclib, Ivacaftor, Bazedoxifene, Niclosamide, and Eltrombopag.

Among these 24 drugs, two FDA-approved drugs drew our attention. First, niclosamide, an antihelminthic drug, exhibited very potent antiviral activity against SARS-CoV-2 (IC_50_ = 0.28 μM). Not surprisingly, its broad-spectrum antiviral effect has been well documented in the literature ^7^ including antiviral properties against SARS- and MERS-CoV ^8,9^. Recently, Gassen et al. demonstrated that niclosamide inhibits SKP2 activity, which enhances autophagy and reduces MERS-CoV replication ^9^. A similar mechanism might be attributable for the inhibition of SARS-CoV-2 infection by niclosamide. Although niclosamide suffers a pharmacokinetic flaw of low adsorption, further development or drug formulation could enable an effective delivery of this drug to the target tissue ^10^.

Second, ciclesonide is another interesting drug candidate for further development although its antiviral potency was much lower (IC_50_ = 4.33 μM) than niclosamide. It is an inhaled corticosteroid used to treat asthma and allergic rhinitis ^11^. A recent report by Matsuyama et al. corroborated our finding of ciclesonide as a potential antiviral drug against SARS-CoV-2 ^12^. A treatment report of three patients who were infected by SARS-CoV-2 in Japan (https://www3.nhk.or.jp/nhkworld/en/news/20200303_20/) warrants further clinical investigation of this drug in patients with COVID-19. Intriguingly, an underlying mechanism for the suppression of viral infection by ciclesonide has been revealed by the isolation of a drug-resistant mutant ^12^. The isolation of the drug-resistant mutant indicated that NSP15, a viral riboendonuclease, is the molecular target of ciclesonide. Together, it is not unreasonable to consider that ciclesonide exhibits a direct-acting antiviral activity in addition to its intrinsic anti-inflammatory function. In the future, siRNA targeting the hormone receptor will allow to assess the extent of direct-acting antiviral activity. With its proven anti-inflammatory activity, ciclesonide may represent as a potent drug which can manifest dual roles (antiviral and anti-inflammatory) for the control of SARS-CoV-2 infection.

Prior to our evaluation of 48 drugs against SARS-CoV-2 infection, we also tested antiviral activity of several other drugs based on the cytopathic effect of the virus in the presence of each drug (Figure 2). In particular, the effect of favipiravir and atazanavir was compared to those of the reference drugs (chloroquine, lopinavir, remdesivir) because favipiravir is considered as a drug candidate for clinical trials and atazanavir was recently predicted as the most potent antiviral drug by AI-inference modeling ^13^. However, in the current work, we did not observe any antiviral activity of either favipiravir or atazanavir.

**Figure 2.**
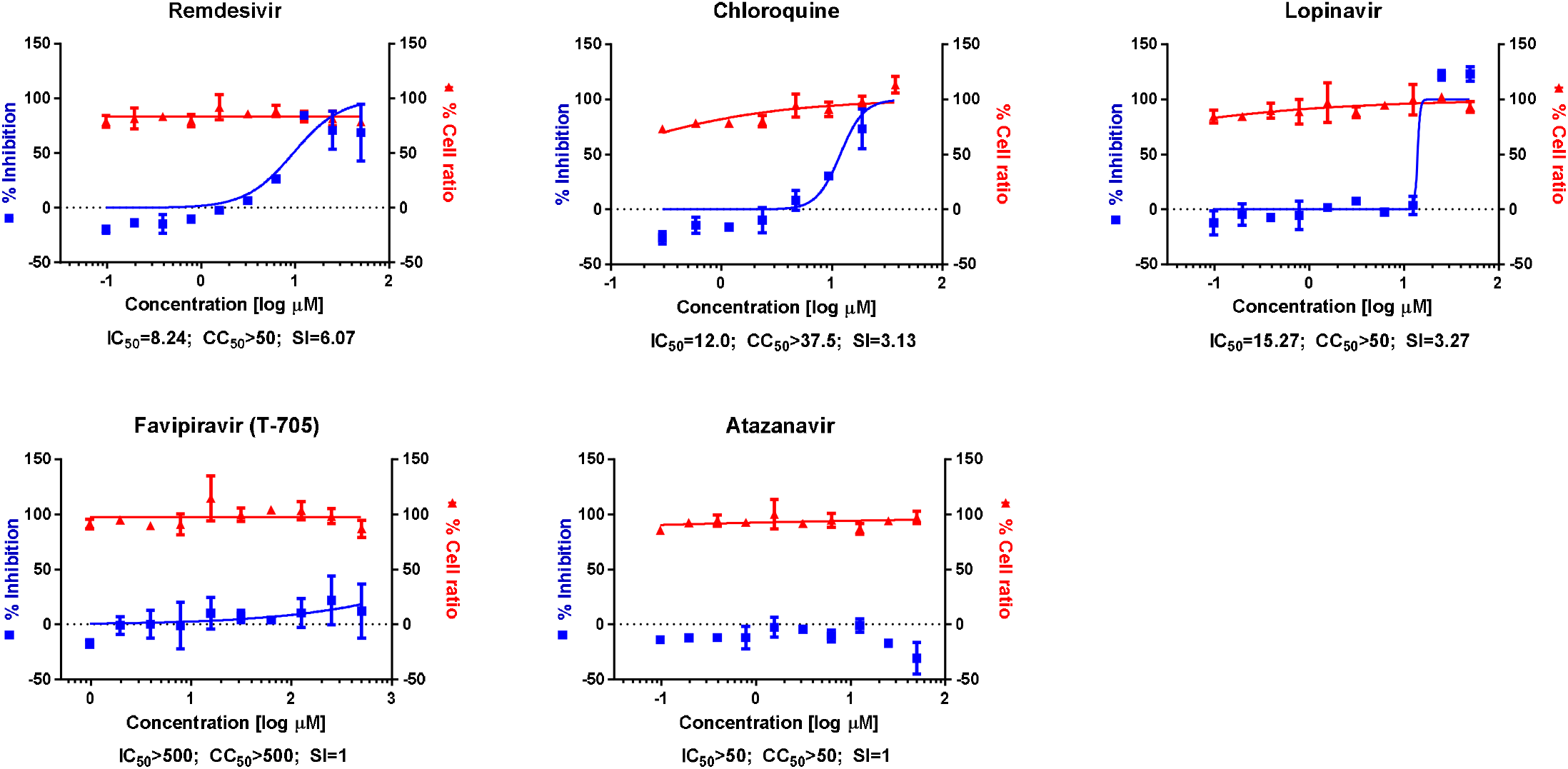
Dose-response curve analysis by cytopathic effect. The blue squares represent inhibition of virus infection (%) and the red triangles represent cell viability (%). Means ± SD were calculated from duplicate experiments.

In summary, we selected and screened 48 FDA-approved drugs based on our SARS-CoV screening and our screening campaign revealed 24 potential antiviral drug candidates against SARS-CoV-2. Our findings could be further validated in an appropriate animal model, and hopefully developed through subsequent clinical trials in order to provide additional therapeutic options for patients with COVID-19.

## Materials and Methods

### Virus and Cells

Vero cells were obtained from the American Type Culture Collection (ATCC CCL-81) and maintained at 37°C with 5% CO_2_ in Dulbecco’s Modified Eagle’s Medium (DMEM; Welgene), supplemented with 10% heat-inactivated fetal bovine serum (FBS) and 1X Antibiotic-Antimycotic solution (Gibco). SARS-CoV-2 (βCoV/KOR/KCDC03/2020) was provided by Korea Centers for Disease Control and Prevention (KCDC), and was propagated in Vero cells. Viral titers were determined by plaque assays in Vero cells. All experiments using SARS-CoV-2 were performed at Institut Pasteur Korea in compliance with the guidelines of the KNIH, using enhanced Biosafety Level 3 (BSL-3) containment procedures in laboratories approved for use by the KCDC.

### Reagents

Chloroquine diphosphate (CQ; C6628) was purchased from Sigma-Aldrich (St. Louis, MO), lopinavir (LPV; S1380) was purchased from SelleckChem (Houston, TX), and remdesivir (HY-104077) was purchased from MedChemExpress (Monmouth Junction, NJ). Chloroquine was dissolved in Dulbecco’s Phosphate-Buffered Saline (DPBS; Welgene), and all other reagents were dissolved in DMSO for the screening. Anti-SARS-CoV-2 N protein antibody was purchased from Sino Biological Inc. (Beijing, China). Alexa Fluor 488 goat anti-rabbit IgG (H + L) secondary antibody and Hoechst 33342 were purchased from Molecular Probes. Paraformaldehyde (PFA) (32% aqueous solution) and normal goat serum were purchased from Electron Microscopy Sciences (Hatfield, PA) and Vector Laboratories, Inc. (Burlingame, CA), respectively.

### Dose-response curve (DRC) analysis by immunofluorescence

Ten-point DRCs were generated for each drug. Vero cells were seeded at 1.2 × 10^4^ cells per well in DMEM, supplemented with 2% FBS and 1X Antibiotic-Antimycotic solution (Gibco) in black, 384-well, μClear plates (Greiner Bio-One), 24 h prior to the experiment. Ten-point DRCs were generated, with compound concentrations ranging from 0.05–50 μM. For viral infection, plates were transferred into the BSL-3 containment facility and SARS-CoV-2 was added at a multiplicity of infection (MOI) of 0.0125. The cells were fixed at 24 hpi with 4% PFA and analyzed by immunofluorescence. The acquired images were analyzed using in-house software to quantify cell numbers and infection ratios, and antiviral activity was normalized to positive (mock) and negative (0.5% DMSO) controls in each assay plate. DRCs were fitted by sigmoidal dose-response models, with the following equation: Y = Bottom + (Top □ Bottom)/(1 + (IC_50_/X)^Hillslope^), using XLfit 4 Software or Prism7. IC_50_ values were calculated from the normalized activity dataset-fitted curves. All IC_50_ and CC_50_ values were measured in duplicate, and the quality of each assay was controlled by Z’-factor and the coefficient of variation in percent (%CV).

### Dose-response curve (DRC) analysis by cytopathic effect (CPE)

Ten-point DRCs were generated for each drug. Vero cells were seeded at 1.2 × 10^4^ cells per well in DMEM, supplemented with 2% FBS and 1X Antibiotic-Antimycotic solution (Gibco) in white, 384-well, μClear plates (Greiner Bio-One), 24 h prior to the experiment. Ten-point DRCs were generated, with compound concentrations ranging from 0.05–50 μM. For viral infection, plates were transferred into the BSL-3 containment facility and SARS-CoV-2 was added at a multiplicity of infection (MOI) of 0.05 and incubated at 37 °C for 72 h. Cell viability was measured using the CellTiter-Glo Luminescent Cell Viability Assay (Promega), according to the manufacturer’s instructions. Antiviral activity was determined by the degree of inhibition of viral cytopathic effect. The results were normalized to positive (mock) and negative (0.5% DMSO) controls in each assay plate. DRCs were fitted by sigmoidal dose-response models, with the following equation: Y = Bottom + (Top □ Bottom)/(1 + (IC_50_/X)^Hillslope^), using XLfit 4 Software or Prism7. IC_50_ values were calculated from the normalized activity dataset-fitted curves. All IC_50_ and CC_50_ values were measured in duplicate, and the quality of each assay was controlled by Z’-factor and the coefficient of variation in percent (%CV).

## Acknowledgements

We thank Drs. Wang-Shick Ryu and Spencer Shorte for their helpful discussion and review of the manuscript. The pathogen resource (NCCP43326) for this study was provided by the National Culture Collection for Pathogens. This work was supported by the National Research Foundation of Korea (NRF) grant funded by the Korean government (MSIT) (NRF-2017M3A9G6068245 and NRF-2020M3E9A1041756).

